# Computational mechanism underlying switching of motor actions

**DOI:** 10.1101/2023.10.27.564490

**Authors:** Shan Zhong, Nader Pouratian, Vassilios Christopoulos

## Abstract

Surviving in a constantly changing environment requires not only the ability to select actions, but also the flexibility to stop and switch actions when necessary. Extensive research has been devoted to understanding how the brain switches actions, yet the computations underlying switching and how it relates to selecting and stopping processes remain elusive. A central question is whether switching is an extension of the stopping process or involves different mechanisms. To address this question, we modeled action regulation tasks with a neurocomputational theory and evaluated its predictions on individuals performing reaches in a dynamic environment. Our findings suggest that, unlike stopping, switching does not necessitate a proactive pause mechanism to delay movement onset. However, switching engages a pause mechanism after movement onset, if the new target location is unknown prior to switch signal. These findings offer a new understanding of the action-switching computations, opening new avenues for future neurophysiological investigations.

## Introduction

Operating effectively in an uncertain and dynamic environment requires not only the ability to accurately prepare and perform actions, but also the flexibility to suppress actions in response to environmental changes. When driving on a city road, we often need to stop at red lights, crosswalks or intersections. However, everyday life rarely calls for complete suppression of actions without subsequent behavioral adjustments. In fact, altering environmental conditions frequently lead us to abandon planned or ongoing actions followed by a switching behavior to adapt to new situations. Understanding the mechanisms of selecting, stopping and switching actions is important for revealing how the brain functions in variable and evolving environments.

The current study aims to dissociate between the mechanisms involved in stopping and switching reaching movements. Normative theories suggest that a current action must first be inhibited by an independent pause mechanism before switching to a new action [1, 2]. This view computationally conceptualizes response inhibition as a race between two “runners” (i.e., processes) - a “go runner” initiated by the presentation of external stimuli and a “stop runner” triggered by a stop signal. If the stop runner wins the competition, the response is inhibited. Otherwise, the response is emitted [2–4]. When switching action is required, the current go process must first be interrupted by the stop process before another go process generates a new action. This theory has received significant support from neurophysiological and functional neuroimaging studies that have identified the basal ganglia (BG), and in particular the subthalamic nucleus (STN), as a key region in canceling an already selected, or currently performed, action when goals change [5–7]. Overall, these studies suggest that switching actions involves the same processes as stopping actions, with the only difference being that a new action is generated after the old one is suppressed.

However, neurophysiological recordings in non-human primates (NHPs) challenged this “go-stop-go” theory, suggesting that an independent inhibitory (i.e., pause) mechanism may not be required when switching actions [8]. Instead, switching actions can be implemented through a competition process between the current and the new action without engaging the stopping circuitry. This theory is considered an extension of the “affordance competition hypothesis” according to which multiple motor actions are formed concurrently and compete over time until one has sufficient evidence to win the competition [9–11]. This hypothesis predicts that action selection is made through a competition within the same circuit that plans and produces the actions themselves. It predicts that the same neurons involved in the initial action selection process remain active in adjusting and even switching between actions during overt behavior [8].

Therefore, the main question is whether the switching process is an extension of the stopping process or it can be implemented through a different mechanism. To address this question, we trained healthy young adult participants to perform reaching movements with a 2D joystick to either a single target or one selected from two targets assigned with different expected rewards. In a subset of trials, participants had to completely stop or switch their reaching movements. To better elucidate the action regulation mechanisms in selecting, stopping and switching of actions, we modeled the reaching tasks within a neurodynamical computational framework that combines dynamic neural field (DNF) theory [12, 13] with stochastic optimal control (SOC) theory [14, 15]. The framework was recently developed to simulate motor behavior and the underpinning neural mechanisms in a variety of visumotor tasks that occur in dynamic and uncertain environments [16,17]. By modeling the experimental tasks within the neurocomputational framework, we provide evidence that reaching planning does not involve a proactive pause mechanism, unless a stop signal is anticipated. Interestingly, the mechanism for switching ongoing actions depends on whether the new target location is known prior to the switch signal. If the participants are not aware of the new target location before they are prompted to correct their reaches, an independent pause mechanism seems to be engaged to suppress the ongoing action. On the other hand, if participants know the new target location prior to the switch signal, switching of action seems to be implemented without engaging the pause mechanism. Overall, our study provide a putative model for the intricate processes involved in stopping and switching actions, opening new avenues to better understand how the brain regulates action in dynamic and uncertain environments.

## Results

### Experimental paradigms

Participants were instructed to perform rapid reaching movements using a 2D joystick in 3 experimental tasks: decision-making (i.e., action selection), stop signal (i.e., outright stopping) and switch task (Fig.1). The decision-making task includes choice trials, during which participants had to decide between two targets (blue circles) associated with different expected rewards that were presented either on the same or opposite visual fields (Fig. 1A). Choice trials were interleaved with instructed trials, in which only one single target was presented in the field. The stop signal task is similar to the decision-making task with the difference that in a random subset of trials (i.e., 33%), the color of the target(s) turned red after a short variable delay (stop signal delay, SSD), indicating that the participants need to immediately stop any planned/ongoing reaching movements. Stop trials occur both in instructed (Fig. 1B) and choice trials (Fig. 1C). If the participants successfully inhibited their action, the SSD increased by 50 ms, making the next stop trial more challenging, otherwise the SSD decreased by 50 ms, making the next stop trial easier. Finally, the switch task is also similar to the decision-making task with the difference that in a random subset of instructed trials (i.e., 33%), the original target was replaced after a short variable delay (switch signal delay, SWSD) by a second target, prompting the participants to switch their actions towards the new target location (Fig. 1D). Similarly, in another 33% random subset of choice trials, the high-reward target disappeared, and the participants had to move to the remaining one (i.e., low-reward target) (Fig. 1E). If the participants successfully switched their actions without crossing the old target location, the SWSD increased by 50 ms, making the next switch trial more challenging. Otherwise, SWSD decreased by 50 ms, making the next switch trial easier.

**Fig. 1.**
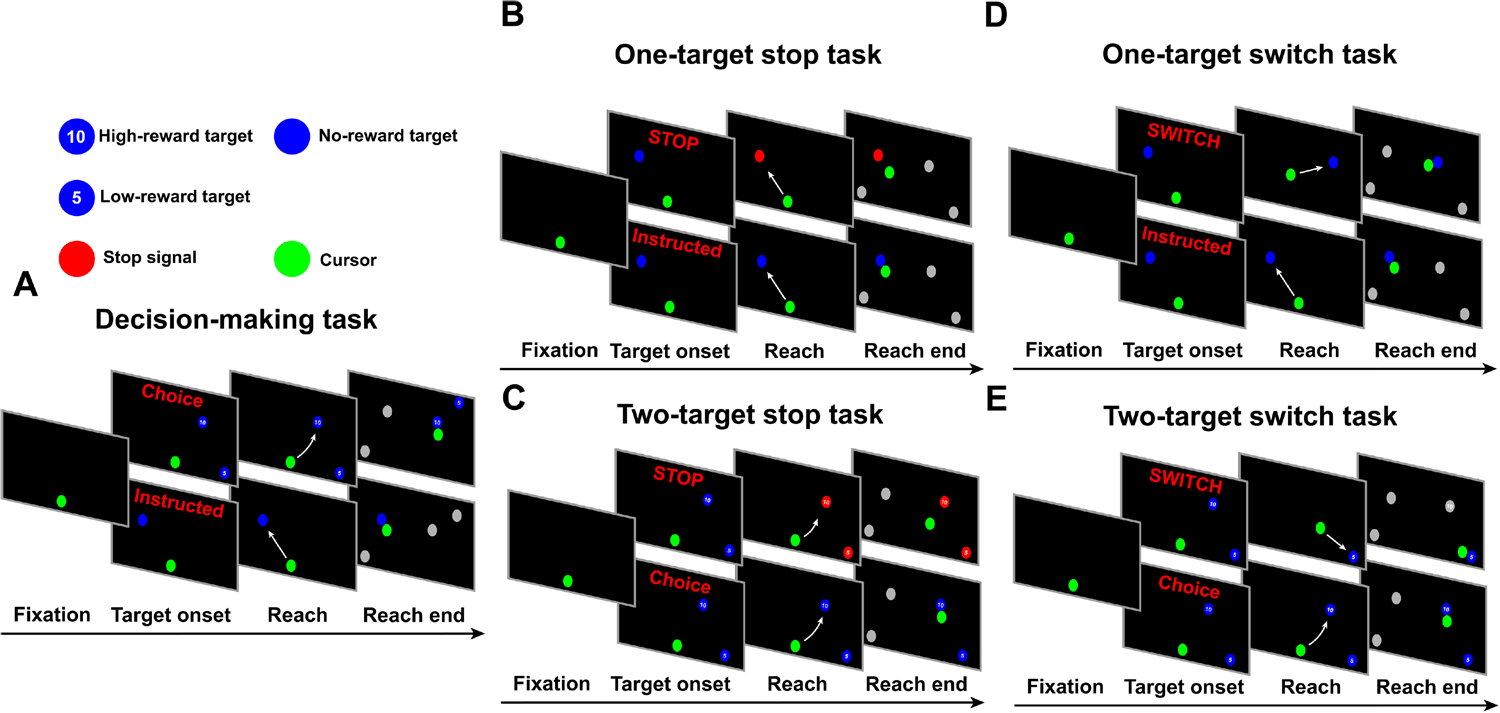
Experimental setup for the decision-making task, the stop signal task and the switch task. (A) Experimental setup for the decision-making task, including instructed trials and choice trials. (B) Experimental setup for the one-target stop task, including stop trials and instructed trials. (C) Experimental setup for the two-target stop task, including stop trials and choice trials. (D) Experimental setup for the one-target switch task, including switch trials and instructed trials. (E) Experimental setup for the two-target switch task, including switch trials and choice trials.

### Motor strategy and performance in selecting, stopping and switching of actions

The participants generated highly stereotyped reaching movements when they were instructed to reach towards a target location (Fig. 2A top panel) or when they were free to choose between two targets presented simultaneously in the screen (Fig. 2A bottom panel). They were also capable of stopping or switching their movements when they were prompted both in instructed (Figs. 2B and C, top panels) and choice trials (Figs. 2B and C, bottom panels). We computed the reaction time (RT) in the instructed and choice trials as an index of motor preparation of the reaching movements. The RT was computed using the time interval between the presentation of the target(s) on the screen and response initiation. Fig. 3A illustrates the average RT across all trials and participants for instructed and choice reaches in the three experimental paradigms. A two-way ANOVA revealed statistical significant differences on RT in the experimental tasks (p < 10*^—^*^6^) and type of movements (i.e., instructed vs. choice) (p < 10*^—^*^6^). A post-hoc multiple-comparison analysis using the Tukey test indicated that instructed trials had shorter RTs than choice trials in all three experimental tasks (p < 10*^—^*^7^). Furthermore, RT was longer both in instructed and choice trials when participants expected a stop signal compared to trials in which they did not anticipate to stop their actions (i.e., decision-making task) (p < 10*^—^*^7^). These findings are consistent with the results from our previous study [17] in which we reported longer RTs when anticipating a stop signal in instructed trials. Notably, we found that participants did not exhibit proactive inhibition in switch tasks, that is, they did not prolong their response when they anticipated to switch their actions both in instructed and choice trials (Fig. 3A). In fact, RT was shorter in the switch task compared to the decision-making task both in instructed and choice trials (p < 10*^—^*^7^), even though participants were given an extra 1.0 s to complete a trial if a switch signal was shown. These results suggest that participants do not take a proactive action for slowing down their movement initiation when a switch signal is anticipated. The shorter RT in the instructed and choice trials of the switch task compared to the decision-making task can reflect an adequate amount of practicing reaching movements - i.e., decision-making task was performed before the switch task.

**Fig. 2.**
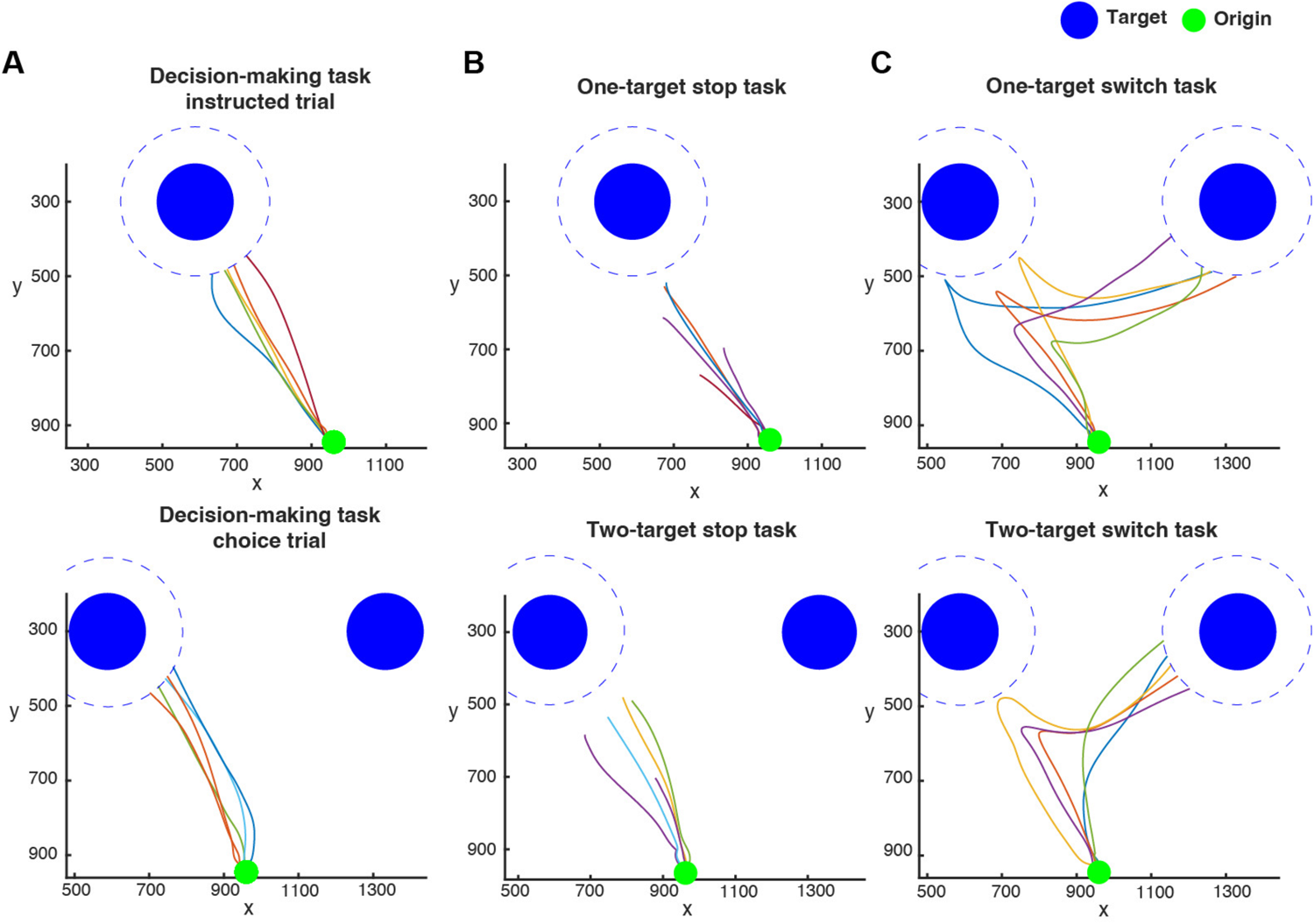
Example human reaching trajectories in the decision-making, stop signal and switch tasks. (A) An example figure that shows 5 example trajectories that starts from the origin (green dot) and go to the instructed target (instructed trial, top) or the selected target (choice trial, bottom). The targets are shown in blue dots. Note that the target would disappear the moment the edge of the cursor touches the edge of the target, and we were tracking the center of the cursor, therefore the trajectories ended a short distance away from the target (marked by blue discontinuous lines). (B) An example figure that shows 5 sample trajectories in the successful stop trials of the one-target stop task (top) and two-target stop task (bottom). Note that the trajectories ended further away from the target than in (A), indicating that that participants did not reach the targets. (C) An example figure that shows 5 sample trajectories in the successful switch trials of the one-target switch task (top) and two-target switch task (bottom). In the case of a one-target switch trial, the target on the left hemisphere is the initial target, which later disappeared and the target on the right hemisphere appears. In the case of a two-target switch trial, the target on the left hemisphere has higher reward, which disappeared shortly after, leaving only the target on the right hemisphere with lower reward.

**Fig. 3.**
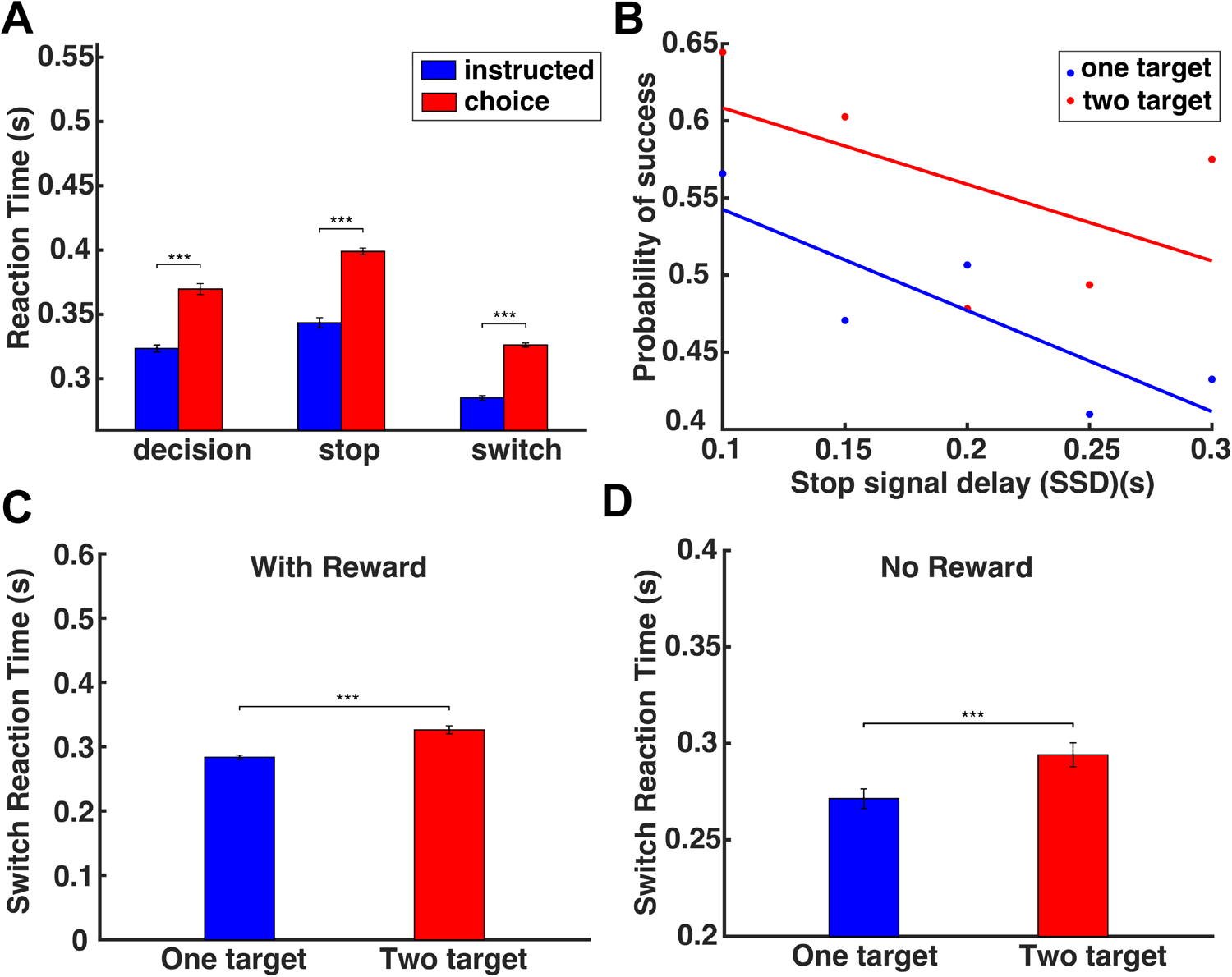
Human performance in the decision-making task, the stop-signal task and the switch task. (A) Bar plots of the RT in the instructed trials (blue) and choice trials (red) of the decision-making task, the stop-signal task and the switch task. Error bars correspond to standard error (SE). (B) The probability to successfully stop an action as a function of the SSD for one-target (blue) and two-target (red) stop tasks. (C) Bar plots of the switch reaction time for the one-target and two-target switch tasks (with reward). (D) Bar plots of the switch reaction time for the one-target and two-target switch tasks (no reward). Error bars correspond to SE.

We also evaluated the performance of the participants by computing the probability to successfully stop an action for different SSD values both in one-target and two-target stop tasks. We found that the probability to stop an action is inversely correlated with SSD - i.e., the longer the SSD, the lower the probability to completely suppress a reaching movement on time - both in one-target and two-target trials (Fig. 3B). Interestingly, the probability of successfully stopping an action was higher when participants were free to choose between two targets (i.e., two-target stop trials) than when they were instructed to reach towards a single target location (i.e., one-target stop trials).

Finally, we computed the time it takes for the participants to respond to a switch signal (switch reaction time, SRT) in instructed and choice trials. The results showed that participants had a shorter switch response when they were instructed to move towards a single target location than when they had to choose between two target locations - i.e., SRT is shorter in one-target switch task than in the two-target switch task (Fig. 3C, two sample t-test analysis, p < 0.001). One potential explanation is that participants became less motivated when they were prompted to switch their reaching movements from a high- to low-reward targets, since motivation is correlated with expected reward [18, 19]. To assess whether the longer SRT is due to the reduction of the expected reward, we recruited 6 participants to perform a modified version of the switch task. The modified switch task was similar to the switch task described above, with the only difference that the targets were not assigned with an expected reward, and the switching occurred once the reach trajectory exceeded a random distance threshold. The results showed that SRT was still longer for two-target switch trials compared to one-target switch trials (Fig. 3D, two sample t-test analysis, p < 0.001). Therefore, we can conclude that people had slower responses to switch their actions when they were aware of the new target location prior to movement initiation (i.e., choice trials) than when they were not aware of the new target location (i.e., instructed trials).

### Predicting motor behavior in action regulation tasks using a neurocomputational theory

To better understand the computations of selecting, stopping and switching actions, we modeled the three experimental tasks within a recently developed neurocomputational theory which models action regulation functions that involve motor inhibition [17]. It combines DNF theory and SOC theory and includes circuitry for perception, expected outcome, effort cost, stop signal, pause mechanism, action planning and execution. It is based on the affordance competition hypothesis, in which multiple actions are formed concurrently and compete over time until one has sufficient evidence to win the competition [9, 20, 21]. In a recent study, we showed that the theory can predict many key aspects of motor behavior in motor tasks that involve selecting and stopping of actions, including spatial characteristics of the reaching trajectories, as well as reaction time, movement velocity, probability of successfully stopping actions, and so on [17]. The architectural organization of the framework is shown in Fig. 4. Each DNF simulates the dynamic evolution of firing rate activity of a network of neurons over a continuous space with local excitation and surround inhibition. The core component of the framework is the “reach planning” field that has two roles: i) activating downstream stochastic optimal controllers that generate actions towards particular directions and ii) integrating information from disparate sources associated with actions, goals, and contextual requirements into a single value (normalized neural activity) that characterizes the relative desirability (i.e., “attractiveness”) of the active actions. The reach planning field receives excitatory input (green arrows) from the “spatial sensory input” field (which encodes the angular representation of the targets in an egocentric reference frame) and “expected outcome” field (which encodes the rewards associated with moving towards particular directions), as well as inhibitory input (red arrows) from the “reach cost” field (encodes the effort required to move towards particular directions) and the “pause” field, which suppresses ongoing (or planned) actions when motor inhibition is required. Each neuron in the reach planning field is connected with a control scheme that generates reaches. Once the activity of a neuron exceeds an action initiation threshold, a decision is made, and the corresponding controller is triggered and generates a sequence of motor actions towards the preferred direction of that neuron. (more details are presented in Materials and Methods section and our recent study [17]).

**Fig. 4.**
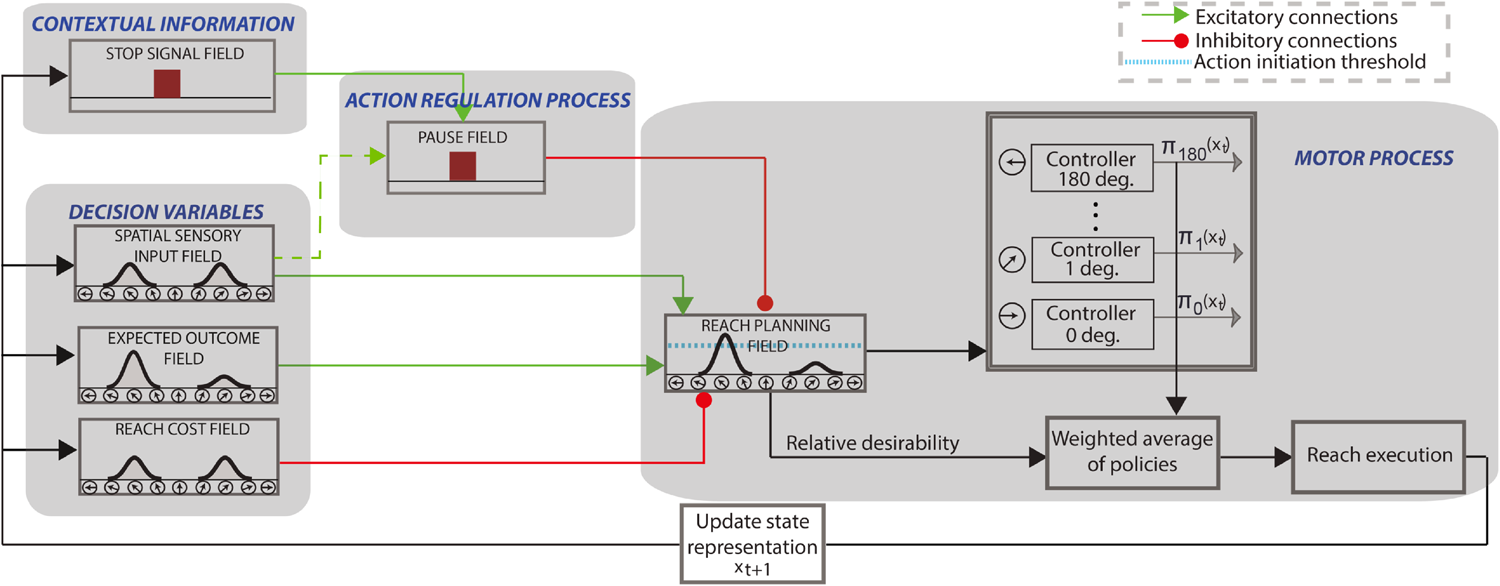
Model Architecture. The reach planning field encodes the planned direction of arm movement in an egocentric reference frame. It integrates the information of sensory information, expected reward and cost, as well as inhibition from the pause field, to generate the “relative desirability”, or the attractiveness of an action policy with respect to alternatives. The relative desirability was used as the weight to compute the final action policy, and a reaching movement is generated, moving the hand towards the finally selected direction.

In the current study, we extended the theory to model the computations involved in switching actions. We considered the following potential architectures that can implement the inhibitory mechanism in action regulation tasks.

- Architecture 1: The pause mechanism is involved in both stopping and switching actions

The pause mechanism is engaged anytime that an action has to be suppressed. In the stop signal task, the pause field is activated during action planning, and further activated to completely suppress all ongoing/planned actions following a stop signal. Similarly, in the switch task, the pause field is also activated during action planning, and further engaged to suppress the ongoing/planned actions before the new action is performed to implement the change of action plan.

- Architecture 2: The pause mechanism is involved only in outright stopping of actions

An alternative architecture suggests that the pause mechanism is involved only in outright stopping of actions. On the contrary, the pause field is not activated during action planning, and switching to new actions is exclusively implemented within the reach planning field – i.e., switching of actions is achieved through a competition within the same circuit that guides the actions themselves. The very similar neurons in the reach planning field that guide action selection will continue to update their activities in the presence of new incoming information to switch the action when needed.

### Modeling reach decisions

We modeled the decision-making task within the neurocomputational framework for both instructed and choice trials. Fig. 5A depicts the simulated neural activity in the reach planning field for typical instructed (top panel) and choice (bottom panel) reaches, respectively. The activity starts at the baseline (resting state) before the target(s) are presented in the field. After the instructed target is presented, the activity of the single neuronal population tuned to the direction of the target increases. Once the activity exceeds the action initiation threshold, a reaching movement is generated towards the target location. In the choice trial, two neuronal populations selective for the targets start competing for selection via mutual inhibitory interactions. This competition leads to a longer RT in movement initiation in choice trials compared to instructed trials. Since the reach planning field receives excitatory inputs from the expected outcome field, the target reward influences choice preferences by shifting the selection bias towards the higher valued target - i.e., the activity of the neuronal population tuned to the higher valued target increases significantly compared to the neural activity associated with the lower valued target. Once the activity of a neuronal population exceeds the action initiation threshold, the competition is resolved and a reaching movement is initiated towards the selected target location.

**Fig. 5.**
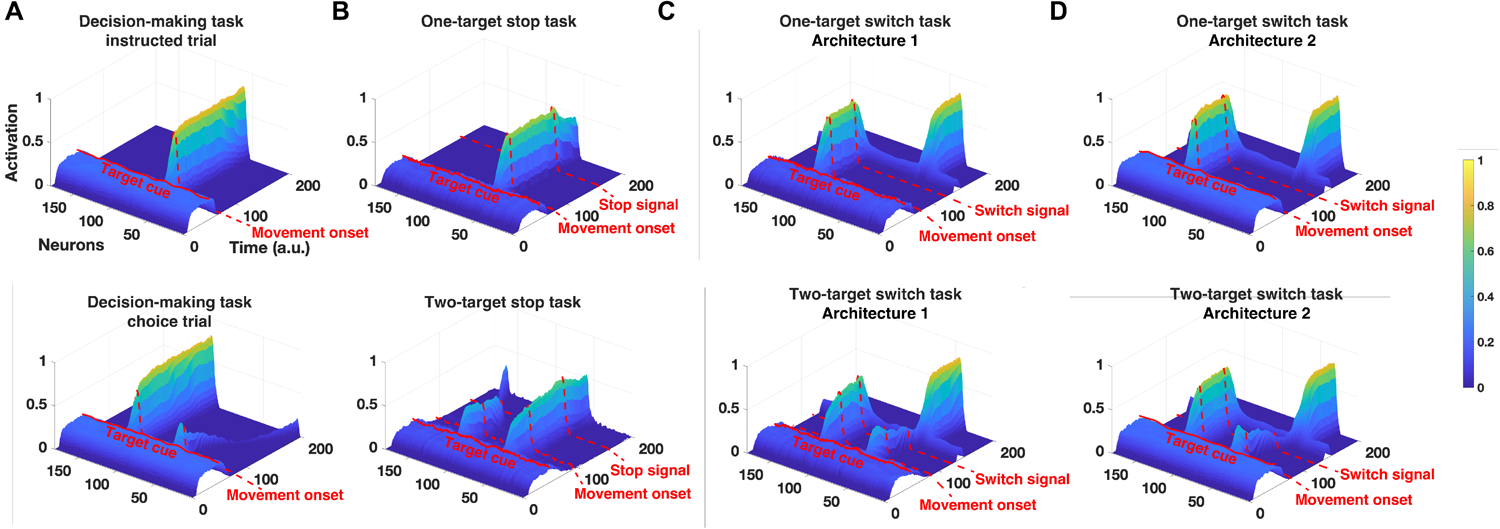
Simulated reach planning field neuronal activity changes in the decision-making task, the stop-signal task and the switch task. (A)-(D): Activity changes of the 181 neurons of the reach planning field during the decision-making task (instructed trial and choice trial)(A), the stop-signal task (one-target stop trials and two-target stop trials)(B), the switch task under architecture 1 (one-target switch trials and two-target switch trials)(C), and the switch task under architecture 2 (one-target switch trials and two-target switch trials)(D).

### Modeling outright stopping of actions

We also modeled the stop signal task within the neurocomputational framework for both one-target and two-target trials. Fig. 5B illustrates the simulated neural activity of the reach planning field for typical instructed (top panel) and choice (bottom panel) reaches that are prompted to completely stop a few time steps after departing from the origin. Note that the pause field is partially activated even before initiating an instructed or a choice reaching movement. This is based on human findings that reaches have longer RTs when participants anticipate a stop signal than when no stop signal is expected (response delay effect, RDE) (Fig. 3, see also [17]). The inhibitory projection from the pause field leads to a reduction in the neuronal signal in the reach planning field when a stop signal is anticipated - i.e., the activity in the reach planning field is lower in the stop trials (Fig. 5B) compared to the instructed and choice trials of the decision-making task (Fig. 5A). Once the stop signal is cued, the activity of the pause field further increases to inhibit the activity of the reach planning field below the action initiation threshold, in order to completely stop the action (see also [17] for more details).

### Modeling switching of actions

Finally, we modeled the switch task within the neurocomputational framework for both one-target and two-target trials, considering two different architectures. Fig. 5C depicts the simulated neural activity of the reach planning field in architecture 1 for typical instructed (top panel) and choice (bottom panel) reaches that are prompted to switch to a new target location a few time steps after departing from the origin. Similar to the stop-signal task, this architecture engages the pause mechanism during the planning phase of the reaching movement, and therefore the activity in the reach planning field is lower when switching is expected than when no switching is anticipated. Once the selected target is replaced by a second target, the activity of the pause field further increases to suppress the current action while the new action is formed towards the new target location. Therefore, switching in this architecture is implemented by the pause mechanism and the mutual inhibitory competition between the current action and the new action. The reach planning field neural activity in the alternative architecture 2 is presented in Fig. 5D. In this scenario, the pause mechanism is not engaged during switching actions. Instead, the switching process is implemented by the same competition process responsible for generating the reaching movement to the initial target location - i.e., the neuronal population tuned to the new target inhibits the neuronal population tuned to the old target. In this case, the reaching neural activity prior to switching action is similar to the activity in tasks where no switching is anticipated.

### Simulated motor behavior in selecting, stopping and switching of actions

We simulated 100 decision-making trials with 50% instructed and 50% choice reaching movements. We also simulated 600 stop signal trials, consisting of an equal split between instructed and choice reaching movements, with 300 trials for each type. Out of these 600 trials, 500 were selected (250 from the instructed category and 250 from the choice category) to be prompted to stop after a short, variable delay (SSD). These 500 trials were diversified using 5 different SSDs, with 50 trials conducted for each SSD. Note that we simulated such a large number of reach trajectories in the stop signal task, so that we explored whether the model can predict the effects of the SSD and the number of targets to the probability of successfully stopping an action. Finally, we simulated 200 trials with 50% instructed and 50% choice reaching movements, but 100 of the trials (50 instructed and 50 choice trials) were prompted to switch the action after a short delay (SWSD). We performed one set of simulations with the architecture 1 (i.e., the pause mechanism is involved in both stopping and switching actions) and another set of simulations with the architecture 2 (i.e., the pause mechanism is involved only in outright stopping of actions). Fig. 6 illustrates a sample of simulated reaching trajectories of the three experimental tasks (consistent across architectures 1 and 2), using only two targets located 60 degrees apart for the sake of simplicity.

**Fig. 6.**
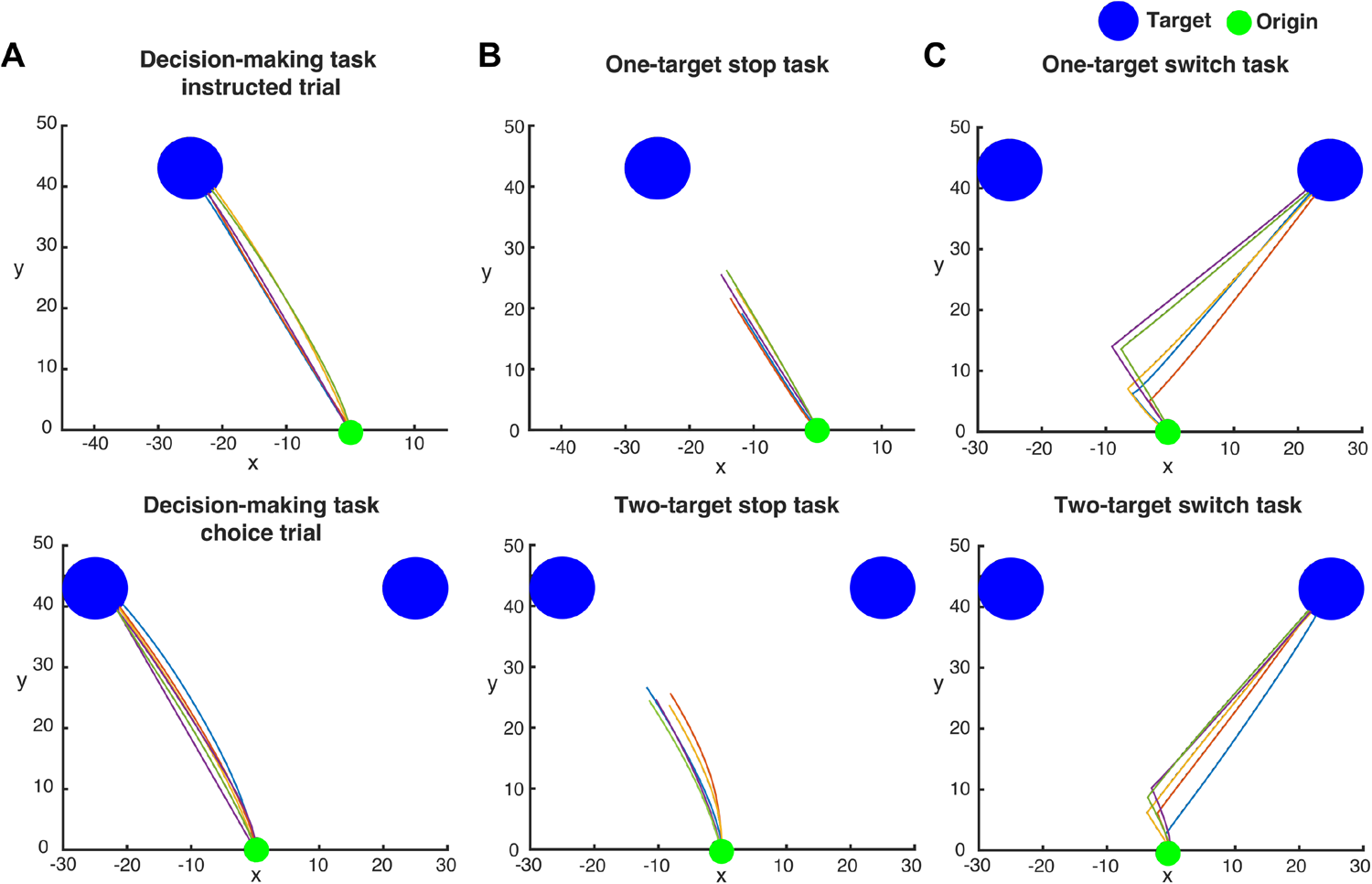
Example simulated reaching trajectories in the decision-making task, the stop-signal task and the switch task. Simulated reaching trajectories that start from the origin (green dot) towards the blue target(s) in the three action regulation tasks. Note that the simulated cursor functioned as a point mass, therefore the trajectories ended at the target location in non-stop trials.

### Reaction time of simulated reaches

Fig. 7A and 7B depict the average RT for the simulated instructed (blue) and choice (red) reaches in the three experimental tasks with (panel A, architecture 1) and without (panel B, architecture 2) the involvement of the pause mechanism in the switch task, respectively. A two-way ANOVA revealed statistical significant differences on RT in the experimental tasks (p < 0.001) and type of movements (i.e., instructed vs. choice) (p < 0.001) for both architectures. A post-hoc multiple-comparisons analysis using the Tukey test showed that choice reaches have a longer RT than instructed reaches (p < 0.001) in all three tasks for both architectures due to the inhibitory action competition when two targets are presented. For the model architecture 1 which involves a pause mechanism for switching actions, no significant differences were observed in RT when anticipating a stop or switch signal, both in instructed (p = 0.959) and choice trials (p = 0.959) (Fig. 7A). On the other hand, for model architecture 2 which does not involve a pause mechanism for switching actions, RT is longer in instructed (p < 0.001) and choice trials (p < 0.001) when a stop signal is anticipated compared to when a switch signal is anticipated (Fig. 7B). Instead, switch trials have approximately the same RT as decision-making trials - i.e., when no switch signal is expected (p = 1.000). The human findings are consistent with the results from the model architecture 2, in which the pause mechanism is not engaged in switching actions during movement planning, since participants responded faster when a switch signal was anticipated than when a stop signal was expected.

**Fig. 7.**
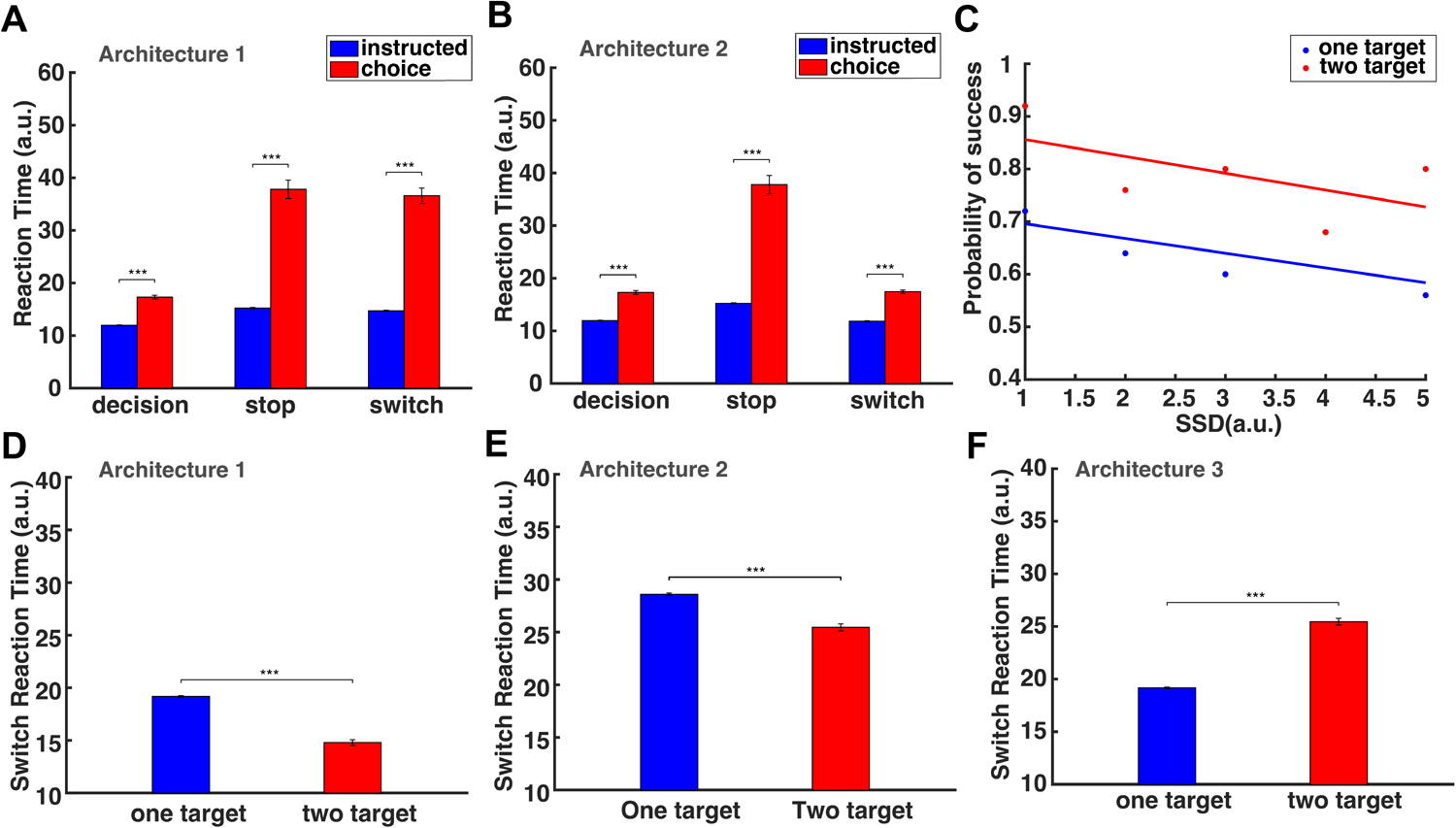
Simulated human behavior in the decision-making task, the stop-signal task and the switch task. (A) Bar plots of the simulated RT in the instructed and choice trials of the decision-making task, the stop-signal task and the switch task (architecture 1). (B) Bar plots of the simulated RT in the instructed and choice trials of the decision-making task, the stop-signal task and the switch task (architecture 2). (C) The simulated probability of successfully stopping a reaching movement as a function of SSD in one-target stop trials (blue) and two-target stop trials (red). (D) Bar plots of the simulated SRT for the one-target and two-target switch trials, when the pause field is involved in the switching process (architecture 1). (E) Bar plots of the simulated SRT for the one-target and two-target switch trials when the pause field is NOT involved in the switching process (architecture 2). (F) Bar plots of the simulated switch reaction time for the one-target and two-target switch trials when the pause field is involved in the switching process in the one-target condition, but not involved in the two-target condition (architecture 3). Error bars correspond to SE.

### Probability to successfully stop actions

We also computed the probability to completely stop a reaching movement in the stop signal task by measuring the number of successful trials at any given SSD in both instructed and choice reaches. Consistent with human findings, the model predicted that the probability to successfully stop reaches is inversely related to SSD - i.e., the higher the SSD, the lower the probability to successfully stop a reaching movement. Notably, and consistent with human findings, the model predicted that the probability to stop an action is higher in the two-target stop trials than in the one-target stop trials for any given SSD (Fig. 7C). The reason is that the activity of the reach neurons in the choice trials is weaker than in the instructed trials (Fig. 5B top and bottom panel) due to inhibitory competition between the neuronal populations that are selective for the two targets. Therefore, the activity of the reach neurons is inhibited faster in the two-target stop trials than in the one-target stop trials, explaining why it is easier to stop an action when you are free to choose between competing targets than when you are instructed to move towards a particular target location.

### Switch reaction time of simulated reaches

Both model architectures predict that the reaction time for switching actions (SRT) will be longer in one-target trials than in two-target trials (Fig. 7D and 7E). The reason is that the reach neural activity associated with the non-selected target in the two-target trials is often not completely suppressed before the switching cue is given. Hence, the activity of the selected target is weaker in the two-target trials as compared to the one-target trials when reaching towards the same target location. Figs. 8A and 8B depict the simulated neural activity of a single reach neuron in an instructed trial (blue trace) and of two reach neurons, one from each population, in a choice trial (red trace) for architectures 1 and 2, respectively. Neurons are centered at the target location(s). Note that the neural activity associated with the new target location exceeds faster the action initiation threshold in the choice trial (red discontinuous trace) than in the instructed trial (blue discontinuous trace) in both architectures. Subsequently, switching of action in choice trials is more readily and quickly made than in instructed trials, because the neural representation of the new target location - the one that was not originally selected - has been formed prior to the switch signal. On the other hand, the neural representation of the new target location has not been formed prior to switch signal in the instructed trials, since the model has been instructed to generate movements towards one single target location without knowing the new target location.

**Fig. 8.**
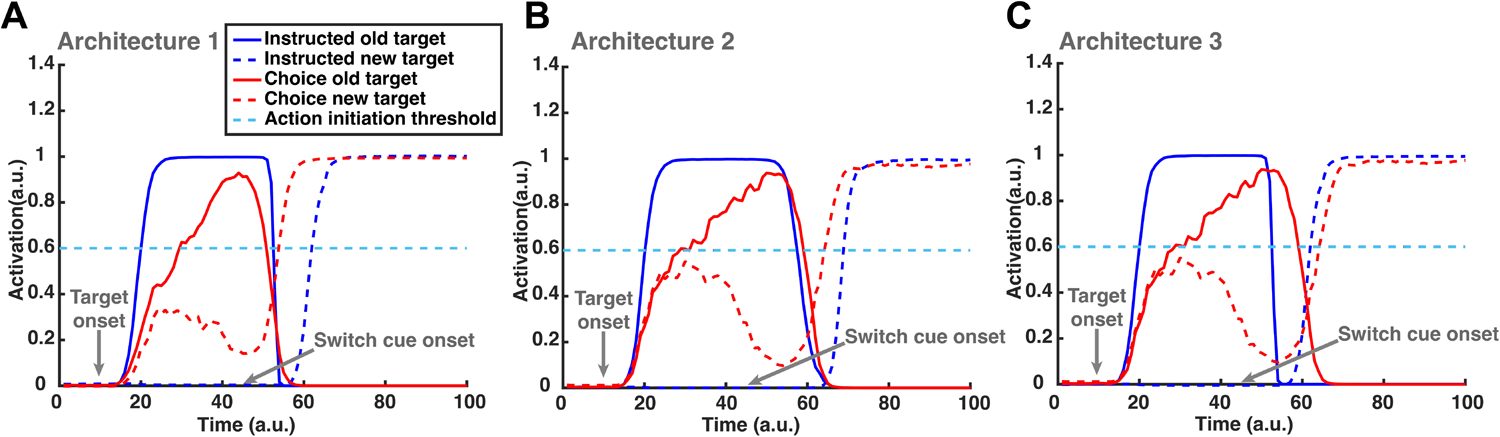
Reach planning field single neuronal activity during switch trials. Simulated reach planning field single neuronal activity changes during one-target and two-target switch trials under architectures 1(A), 2(B) and 3(C). Color code: Blue: one-target switch trial. Red: two-target switch trial.

Interestingly, model predictions contradict the human findings. Participants exhibited longer SRT in two-target switch trials than in one-target switch trials, both when they performed the reward-based and the non-reward-based (i.e., internally-guided) reaching tasks. One potential explanation is that one-target switch trials are governed by different switching mechanisms than the two-target trials during action execution. For instance, an alternative third architecture is that the “pause mechanism” is activated only in the instructed trials to inhibit the ongoing action. The reason is that the neural activity of the current action is too strong to be inhibited on-time by the new incoming action, which will be formed only after the switch signal. On the other hand, the neural activity of the selected target in the two-target trials is weaker compared to the one-target trials, due to the residual activity from the unselected target that inhibits the neural activity of the selected target. Therefore, the new incoming action, which is already formed before the switch signal, can inhibit the current action when needed without the contribution of the pause mechanism. Fig. 8C illustrates the simulated neural activity of a single reach neuron in an instructed trial (blue trace) and of two reach neurons, one from each population, in a choice trial (red trace) under this model architecture. We simulated the switch task within the new architecture and found that the instructed trials have shorter SRT than the choice trials (Fig. 7F). Taken together, our results suggest a third potential architecture (i.e., a combination of the architectures 1 and 2) of the switching process, in which the pause mechanism is not active during action planning, but it is activated in action execution once a switch signal is detected, but only when reaches are instructed to a single target location.

## Discussion

### General

Humans can rapidly regulate actions according to evolving environmental demands. At the same time, impairments of action regulation have been identified across a number of neurological and psychiatric diseases, including Parkinson’s disease (PD), obsessive compulsive disorder (OCD), and Tourette syndrome [22–29]. Given the ubiquity of action regulation in everyday life, its critical role in survival, and its impairment across a variety of neurological and psychiatric diseases, understanding the mechanism of action regulation is of high value and impact. A key component of action regulation is action inhibition that occurs when stopping unwanted or inappropriate actions. Normative theories have also suggested that action inhibition plays a critical role in switching between actions in response to environmental changes. An ongoing (or planned) action has to be first inhibited, before switching to a new action [30, 31]. A popular view is that when the pre-supplementary area (pre-SMA) detects the co-activation of different responses - a current response and a new response for switching action - it activates the STN to temporarily suppress the current response (reviewed by [32, 33]). However, an alternative theory suggests that switching action might not always necessitate an independent action inhibition process (i.e., a pause mechanism). Instead, the same neurons within the motor areas that are involved in selecting the initial action will continue to be involved in adjusting and even switching actions during overt behavior [8, 21]. This suggests that the brain may not always need to engage an independent inhibition mechanism to switch actions, particularly in scenarios where multiple potential actions are concurrently represented. Therefore, the mechanisms underlying how the brain selects, stops and switches between actions, as well as how these seemingly disparate functions inter-relate both behaviorally and computationally still remain elusive.

Our study focuses on the intricate interplay between action selection, stopping, and switching, and how these processes are interrelated within the human brain. We trained healthy young adult participants to perform reaching movements with a 2D joystick for running behavioral tasks that involve action selection, stopping and switching under conditions that one or two target(s) were presented at the beginning of each trial. Our findings show that when participants anticipate a stop signal, they delay initiating a movement in order to increase the chance to successfully stop the action. Interestingly, the results did not reveal this proactive inhibitory behavior when participants anticipate a switch signal, both when they were instructed to move towards a target location or when they were free to choose between two targets. Instead, they initiated faster movements similar to trials in which no adjustment in behavior was required - i.e., decision-making task without stop or switch signals. These findings contradict the normative theory - at least during the planning phase of reaching movements - according to which switching behavior is an extension of stopping behavior with a new action to generate after the current one is suppressed through a pause mechanism. Interestingly, participants were faster to switch their actions in instructed than in choice trials, even though the new target locations were known in choice trials before the participants were prompted to correct their actions and move towards the new targets. Additionally, participants had better performance to completely stop an action in choice trials than in instructed trials regardless of when the stop signal was cued.

By modeling this motor behavior within a recently developed neurocomputational theory [17], we predicted that the pause mechanism is engaged only when an outright stopping of action is anticipated. However, this action regulation mechanism fails to explain the faster responses to switch *ongoing* actions in instructed trials compared to choice trials, suggesting there might be a different mechanism for switching ongoing actions between instructed and choice movements. We considered an alternative hypothesis in which the pause mechanism is engaged to switch actions only in instructed trials. On the other hand, switching action in choice trials is implemented through a competition between the ongoing and the new actions. The rationale behind this hypothesis is that the neural activity associated with the new action has not been formed prior to the switch signal in the instructed trials. Therefore, there might not be enough time for the new action to form and suppress the ongoing action without engaging a pause mechanism that further assists in the suppression process. On the other hand, the new action (i.e., the action associated with the unselected target) has already been formed in the choice trials prior to the switch signal and therefore switching can be implemented on-time through the competition between the ongoing and the new action. This mechanism generates reaching movements with longer SRT in choice trials compared to instructed trials. Overall, this study advances our understanding of the action regulation mechanism, providing evidence that the involvement of the pause mechanism is not constant but rather selectively activated during specific phases of action switching.

### Stopping actions and affordance competition hypothesis

One of the important findings in our study is that participants performed better for stopping reaching movements when they were free to choose between competing targets than when they were instructed to move towards a target location regardless of the SSD. This finding is in favor of the affordance competition hypothesis, according to which multiple actions are formed concurrently and compete over time until one has sufficient evidence to win the competition [20]. Even when a decision is made and the reaching trajectory aims towards the selected target location, there is residual neural activity associated with the unselected target that suppresses the neural activity of the selected target (see Fig. 5B bottom panel). On the other hand, the lack of competition in the instructed trials make the neural activity, which is selective for the selected target location stronger than the neural activity that is selective for the selected target location in the choice trials. Therefore, it is easier to inhibit the motor activity below the action initiation threshold and stop a reaching movement in the choice trials than in instructed trials, explaining why participants performed better for stopping actions whey they were free to choose between two targets than when they were instructed to move towards a particular target.

### Switching actions

One of the main findings in our study is that people delayed to initiate an action when expecting a stop signal both in instructed and choice trials. This RDE effect has been reported in previous studies [17, 34, 35] and has been associated with an “active braking mechanism” that increases the chance of abandoning a response in case stopping is required [36]. Interestingly, RDE was not found in the switch trials - i.e., participants did not proactively slow down their response when a switch signal was anticipated. This suggests that the pause mechanism is not engaged when switching of action is anticipated. However, when we modeled the switch task without the involvement of the pause mechanism, we predicted that it takes longer to switch a reaching movement in instructed than in choice trials. This is against the human findings, in which instructed trials have shorter SRT than choice trials. To account for these results, we modeled the mechanism of switching action within a new architecture, in which the pause mechanism is engaged only in switching instructed trials. In this architecture, two inhibitory mechanisms are involved in switching actions: a) the pause mechanism that suppresses the current action and b) the inhibitory competition between the current and the new action. On the other hand, reaching movements in choice trials switch direction only through the inhibitory competition between the current action and the new action. Therefore, a reasonable question is if instructed trials require a pause mechanism for switching ongoing reaches, why people did not exhibit a proactive planning behavior when they anticipated a switch signal, as they did when they anticipated a stop signal. One potential explanation is that the pause mechanism is not activated prior to movement initiation, since the goal is not to completely stop the ongoing action, but to reduce the neural activity associated with that action while the activity of the new one is formed. Another explanation is that switching of actions already involves an inhibitory mechanism - i.e., the inhibitory competition between the current and the new action. Therefore, the pause mechanism acts as a auxiliary mechanism to inhibit on-time the current action while the new one is formed. Another scenario that could account for the different SRTs between instructed and choice reaches is that when people decide between two targets, the decision process continues even after movement initiation as has been reported in “change-of-mind” studies [37, 38] - i.e., initial decisions are revised after movement onset. Once the switch signal is cued, in the two-target switch trials, people might need to first stop the decision-making process and then re-direct their movements towards the new target location, which is an overall slower process than when they need only to re-direct their movements (i.e., one-target switch trials).

### Building hypotheses for future experiments

We need to point out that the neurocomputational theory used in this study is a systems-level theory that aims to qualitatively predicts the motor behavior, as well as the underpinning neural mechanisms in the three experimental tasks. It can predict many key features of motor behavior in action regulation tasks that involve action inhibition. However, it exhibits the same limitations as any computational theory on what it can predict and whether it can dissociate between alternative and competing hypotheses. For instance, it predicts a particular mechanism of action regulation that can explain the longer SRT that participants exhibited in choice than in instructed trials. However, this does not exclude alternative scenarios, such as the slow process for suppressing first an evolving decision-making process before switching action, that can also generate motor behaviors with longer SRT in choice than in instructed trials. However, this limitation can also be viewed as an advantage of the neurocomputational theory, since it generates theory-driven hypotheses that can be tested in future experimental studies. For instance, the involvement or not of a pause mechanism can be tested by measuring activity from STN in humans and/or animals that perform motor tasks that require switching of actions.

## Conclusion

Overall, our study aims to better elucidate the computations underlying switching of actions and to assess whether switching is an extension of the stopping process, or it involves a different mechanism. To do so, we trained people to perform reaching movements in dynamic environments that involve stopping and switching of actions, and modeled their motor behavior within a neurocomputational framework. The results showed that action planning involves a pause mechanism only when a stop signal is anticipated. How-ever, when a switch signal is anticipated, the pause mechanism is not engaged to delay movement initiation. Interestingly, the results suggest different mechanisms for switching actions when people are instructed to move towards a single target, and when they are free to choose between two targets. To conclude, our study provides novel insights into the computations of action regulations that involve action inhibition, open new doors for further investigation of the action regulation mechanisms in neurophysiological studies.

## Materials and methods

### Ethics Statement

The study was approved by the University of California, Riverside Review Board and all individuals signed a written informed consent before participating.

### Participants

A total number of 20 neurologically healthy adults (9 females) participated in the study. The ages at the time of the experiment were 23.98 ± 4.88 (mean ± SD) years old.

### Stimuli and experimental procedure

#### General

All experiments were programmed using Psychophysics Toolbox Version 3 (PTB-3) for Matlab. Experimental setup is illustrated in Fig. 1. The participants sat in an experimental room approximately 60 cm from an LED monitor (dell P2419HC). A two-dimensional joystick (Thrustmaster T.16000M FCS) was positioned in front of the sitting participants, with the base at the level of their elbows. The real-time position of the joystick was presented on the screen by a green circular cursor (⇠5.5 cm diameter). Participants were familiarized with the task by running a set of training trials, including reaches to one (instructed) and two (choice) target trials, both with and without stop and switch cues. Once they felt ready and comfortable with the tasks, the actual experiment started.

#### Decision-Making Task

In the decision-making task, participants were free to choose between two targets by moving the cursor towards the selected target location. Choice trials were randomly interleaved with instructed trials, during which the participants had to reach toward a single target location (50% instructed and 50% choice trials). A trial started with the cursor appearing at the bottom center of the screen. After 1.0-1.1 s, a 5.5 cm diameter circular blue cue (instructed trial) or two blue cues (choice trial) were presented on the screen, indicating the location of the target(s). The targets were displayed at four possible locations on the screen, each positioned 20.37 cm from the joystick’s starting point. The targets were placed at angles of 0, 60, 120, and 180 degrees, making them equidistant and 60 degrees apart from one another. The participants were instructed not to move the joystick before the target(s) appear on the screen. Once the target(s) were presented, the participants had to move the cursor towards the instructed or to the chosen target location within 1.0 s. In the choice trials, the two targets were marked with a different number (5 or 10) that reflects the reward value. Although the two targets were assigned with different reward values, both options were regarded as “correct”. A reaching movement was considered successful if the cursor touched the target within 1.0 s after the presentation of the target(s). If the participants failed to reach to the target within 1.0 s or initiate a movement prior to target(s) presentation, the trial was aborted. After each trial, the participant had to move the cursor back to the original starting position. Otherwise, they received a warning signal “please move the joystick back to the center to start the next trial”. The participants performed 2 blocks with 48 trials each (2 blocks x 48 trials = 96 total trials).

#### Stop signal task

The stop signal task is similar to the decision-making task with the difference that participants had to completely stop their actions in a random subset of trials (33%). It is consisted of one-target stop task (i.e., instructed trials with stop signal) and two-target switch task (i.e., choice trials with stop signal) that were performed in separate blocks of trials. In the stop trials, the color of the target(s) turned red after a short delay (stop signal delay, SSD), signifying the immediate need to abandon the action. The participants were informed that stopping and reaching to target(s) are equally important. A trial started with the joystick cursor presented at the bottom of the screen. After 1.0-1.1 s, a single blue cue (instructed trial) or two blue cues associated with different reward values (choice trial) were presented on the screen, and the participants had to initiate a movement towards either the single target or the selected target location within 1.0 s. If the target(s) turned red, the participants had to abandon the movement immediately. If participants successfully inhibited their actions in a timely manner, the target(s) stayed on the screen for the remaining 1.0 s, and the trial was considered successful. Then, the SSD increased by 50 ms, making the next stop trial more challenging. If participants failed to inhibit their actions, the target and the cursor disappeared, and the screen turned black for the remaining of the 1.0 s plus 0.5 s. In this case, the SSD decreased by 50 ms, making the next stop trial easier. The participants performed 2 blocks of the one-target stop task - each block comprised 60 trials, of which 20 were stop trials (2 blocks x 60 trials = 120 total trials). They also performed 2 blocks of the two-target stop task - each block comprised of 72 trials, of which 24 were stop trials (2 blocks x 72 trials = 144 total trials).

#### Switch task in reward-based decisions

The switch task was also similar to the stop signal task with the difference that participants had to perform corrected movements, instead of completely stopping their actions. It consists of one-target switch task (i.e., instructed trials with switch signal) and two-target switch task (i.e., choice trials with switch signal) that were performed in separate blocks of trials. In both tasks, the switch trials constitute a random 33 % of the total trials. In one-target switch trials, the target was replaced after a short variable delay (named switch signal delay, SWSD) by a second target at a new location, prompting the participants to switch their actions towards the new target location. In two-target switch trials, the high-reward target was removed after SWSD, and the participants had to correct their actions by moving towards the remaining target (low-reward target). The participants were informed that switching and reaching are equally important. A trial started with the joystick cursor presented at the bottom of the screen. After 1.0-1.1 s, a single blue cue (instructed trial) or two blue cues associated with different reward values (choice trial) were presented on the screen, and the participants had to initiate a movement towards either the single target or the selected target location within 1.0 s. If switching action was prompted, an extra 1.0 s was given to the participants to complete their actions. If the participants successfully switched to the new target without crossing the location of the old target, the trial was considered successful, the screen turned black and a new trial started. In this case, the SWSD increased by 50 ms, making the next switch trial more challenging. If the participants failed to switch to the new target (crossing the location of the old target or failed to arrive at the new target location), the trial was aborted and the SWSD decreased by 50 ms, making the next switch trial easier. The participants performed 2 blocks of one-target switch tasks with 60 trials in each block - 40 instructed trials without switching and 20 one-target switch trials (2 blocks x 60 trials = 120 total trials). They also performed 2 blocks of two-target switch tasks with 72 trials in each block - 48 choice trials without switching and 24 two-target switch trials (2 blocks x 72 trials = 144 total trials).

#### Switch task in internally-guided decisions

We are also interested in exploring whether the reward value affects the switching behavior. To do so, we recruited 6 participants to perform the switch task without assigning reward values to the targets. Instead, the participants performed “internally guided decisions” - free choices that are not informed by any external contingencies. The switch signal was cued based on the traveled distance from the origin - i.e., “distance-threshold”. The distance-threshold was set to a random number between 1.2% to 3.2% of the total distance between the origin and the target(s) location. The switch task without reward also consists of one-target switch task and two-target switch task that were performed in separate blocks of trials. In one-target switch trials, the original target was replaced by a new target when the cursor exceeded the distance-threshold, prompting the participants to correct their actions towards the new target location. In two-target switch trials, the target that was closer to the cursor disappeared, prompting the participants to move towards the remaining target. Everything else was exactly the same with the switch task in reward-based decisions. The participants performed 5 blocks of one-target switch task with 60 trials in each block - 40 instructed trials without switching and 20 one-target switch trials (5 blocks x 60 trials = 300 total trials). They also performed 5 blocks of two-target switch task with 72 trials in each block - 48 choice trials without switching and 24 two-target switch trials (5 blocks x 72 trials = 360 total trials).

### Statistical Analysis

Cubic smoothing spline interpolation was used to smooth the joystick movement trajectories and to compute the velocity of the movements. RT was defined as the time between the target appearance and the time that movement velocity exceeded 10% of the maximum velocity within the trial. RTs faster than 100 ms were excluded from further analysis because anticipation is considered to be involved before participants initiate an action. RT outliers (RTs more than 3 standard deviations from the mean RT) were also excluded. RTs across all participants were pooled together, and two-way ANOVA analyses were performed to determine the group differences in RTs. We also computed the reaction time for switching reaching movements (i.e., SRT) as the time between the switch signal (i.e., instructed/selected target disappears) and the time when the joystick starts to move towards the new target location. A two sample t-test was performed to compare the differences in SRT.

### Computational framework

We utilized a neurodynamical computational framework that was recently proposed by our group to model action regulation tasks that involve motor inhibition, such as selecting, stopping, and switching actions [17]. The computational framework is based on DNF theory and SOC theory, and uses a series of DNFs to model the neuronal circuitry for perception, expected outcome, effort cost, context signal, action planning, and execution. The functional properties of each DNF are determined by the lateral inhibition within the field and the connections with other fields in the architecture. The projections between the fields can be topologically organized – i.e., each neuron i in the field drives the activation at the corresponding neuron i in the other field, or unordered – i.e., each neuron in one field is connected with all neurons in the other field.

The architectural organization of the framework is shown in Fig.4. The DNF platforms (except for the stop signal field and the pause field) consist of 181 neurons, with preferred direction between 0 and 180 degrees. The “spatial sensory input” field encodes the angular representation of the target(s), and the “expected outcome” field encodes the expected reward for reaching to a particular direction. The outputs of these two fields send excitatory projections (green arrows) to the “reach planning” field in a topological manner. The “reach cost” field encodes the effort cost required to implement a reaching movement at a given time and state. It sends inhibitory projections (red arrow) to the reach planning field to penalize high-effort actions. For instance, an action that requires changing of moving direction is more “costly” than an action of keeping going in the same direction. The “stop signal” field consists of 100 neurons and is activated when a stop cue signal (i.e., the color of the target turned red) is detected. It is linked via one-to-all excitatory projections with the pause field. The “pause field” is linked via one-to-all inhibitory connections with the reach planning field. Once a stop cue is detected, the pause field quickly suppresses the activity of the reach planning field to stop the planned or ongoing actions.

The normalized activity of the reach planning field describes the relative desirability d*_i_*(t) of each “reach neuron” with respect to the alternative neurons at time t – i.e., the higher the activity of a reach neuron i, the higher the desirability to move towards the preferred direction *_i_* of this neuron with respect to the alternatives at a given time t. Each neuron i in the reach planning field is connected with an optimal control scheme that generates reaches. Once the activity of a particular neuron i exceeds an “action initiation threshold”, the controller is triggered and generates an optimal policy ⇡*_i_* – i.e., a sequence of motor actions towards the preferred direction of the neuron i. Hence, a decision is made once a neuronal population exceeds the action initiation threshold and the performed action ⇡*_mix_* (x*_t_*) is given as a mixture of the active policies (i.e., policies with active reach neurons) weighted by relative desirability values of the corresponding neurons at any given time and state:

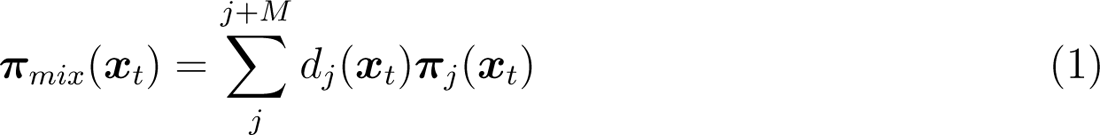

where x*_t_* is the state of the system at time t (i.e., position, velocity, acceleration, etc.), d*_i_*(t) is the normalized activity of the neuron i, and ⇡*_i_* is the optimal policy generated by the controller connected with neuron i. Because desirability is time- and state-dependent, the weighted mixture of the individual policies ⇡*_mix_*(x*_t_*) can be reprogrammed based on the new incoming information. For more details about the mathematics underlying the computational framework, see [16, 17].

## Acknowledgments

Research reported in this publication was supported by National Institute of Neurological Disorders and Stroke under award number U01NS132788. The content is solely the responsibility of the authors and does not necessarily represent the official views of the National Institutes of Health.

## Author Contributions

V.C. and N.P. conceived the study; V.C. and N.P. designed the experiment; S.Z. recruited subjects and collected the data; S.Z. performed the data analysis; S.Z. designed the neurocomputational model and performed the simulations; S.Z. drafted the manuscript with substantial contribution from V.C.; S.Z., N.P. and V.C. revised and approved the manuscript.

## Data Availability

All relevant data within the manuscript and its supporting information files are available under: https://osf.io/cgfaj/.

---

| Model Parameters |  |  |
| --- | --- | --- |
| Parameters | Description | Value |
| $\eta_{loc}$ | Visual input gain | 8.5 |
| $\eta_{reward}$ | Expected outcome input gain | 2.5 |
| $\eta_{cost}$ | Action cost input gain | -0.1 |
| $\eta_{pau}$ | Pause input gain | -4.0 |
| $\gamma$ | Action initiation threshold | 0.6 |

| Spatial sensory input field & Expected outcome field parameters |  |  |
| --- | --- | --- |
| Parameters | Description | Value |
| $\tau$ | Time constant | 5.0 |
| $c_{exc}$ | Amplitude of excitatory portion of weight kernel | 0 |
| $c_{inh}$ | Amplitude of inhibitory portion of weight kernel | 0 |
| $\sigma_{exc}$ | Width of excitatory portion of weight kernel | 5.0 |
| $\sigma_{inh}$ | Width of inhibitory portion of weight kernel | 40.0 |
| $h$ | Resting activity level | -5.0 |
| $q$ | Noise level | 0.25 |
| $\sigma_q$ | Width of noise kernel | 5.0 |
| $\beta$ | Steepness of sigmoid activity function | 1.0 |
| $g$ | Strength of global inhibition | 0.8 |

| Reach planning field parameters |  |  |
| --- | --- | --- |
| Parameters | Description | Value |
| $\tau$ | Time constant | 5.0 |
| $c_{exc}$ | Amplitude of excitatory portion of weight kernel | 20 |
| $c_{inh}$ | Amplitude of inhibitory portion of weight kernel | 5 |
| $\sigma_{exc}$ | Width of excitatory portion of weight kernel | 5.0 |
| $\sigma_{inh}$ | Width of inhibitory portion of weight kernel | 180 |
| $h$ | Resting activity level | -5.0 |
| $q$ | Noise level | 0.5 |
| $\sigma_q$ | Width of noise kernel | 5.0 |
| $\beta$ | Steepness of sigmoid activity function | 1.0 |
| $g$ | Strength of global inhibition | 1 |

| Pause field parameters |  |  |
| --- | --- | --- |
| Parameters | Description | Value |
| $\tau$ | Time constant | 5.0 |
| $c_{exc}$ | Amplitude of excitatory portion of weight kernel | 0 |
| $c_{inh}$ | Amplitude of inhibitory portion of weight kernel | 0 |
| $\sigma_{exc}$ | Width of excitatory portion of weight kernel | 5.0 |
| $\sigma_{inh}$ | Width of inhibitory portion of weight kernel | 25.0 |
| $h$ | Resting activity level | -5.0 |
| $q$ | Noise level | 0.25 |
| $\sigma_q$ | Width of noise kernel | 5.0 |
| $\beta$ | Steepness of sigmoid activity function | 1.0 |
| $g$ | Strength of global inhibition | 0 |

